## Supplemental Tables for "A genetic, genomic, and computational resource for exploring neural circuit function": TableS5.docx

| **Genotype** | **Figure(s)** | **Labeling** |
| --- | --- | --- |
| 20XUAS-CsChrimson-mVenus in attP18/ w^1118^; split-GAL4-AD/+; split-GAL4-DBD/+ | 1D, S1A (SS00116, SS00090, SS00078) | anti-GFP, anti-Brp  DPX mounting  20x objective  Maximum Intensity projections through entire brain |
| pJFRC19-13XLexAop2-IVS-myr::GFP in su(Hw)attP8, pJFRC21-10XUAS-IVS-mCD8::RFP in attP3/ w^1118^ ; split-GAL4-AD/+; split-GAL4-DBD/ UAS unc84 2xGFP | 1D, S1A, S2A (ninaEfl-GAL4, SS55442, SS55437, SS55438, SS08909, SS55440, SS55439, SS55441) | anti-dsRed (recognizes RFP), anti-GFP (not shown), anti-CadN  Slowfade mounting  Maximum Intensity projections through entire brain (1D, S1A) (20x objective)  Single confocal sections (S2A) (40x objective) |
| Published images from Aso et al 2015; image stacks downloaded from https://www.janelia.org/split-GAL4 | 1D (MB312B, MB419B) | see Aso et al 2015  Maximum Intensity projections through entire brain |
| w^1118^; split-GAL4-AD/+; split-GAL4-DBD/pJFRC51-3XUAS-IVS-Syt::smHA in su(Hw)attP1, pJFRC225-5XUAS-IVS-myr::smFLAG in VK00005 | 1D, S1A, S1C, C’, S2A, S2B (all images except lines mentioned above and single cell images) | anti-FLAG, anti-HA (not shown), anti-Brp  DPX mounting  resampled views generated with Vaa3D ( 63x objective)  Maximum Intensity projections through entire brain (20x objective) |
| w^1118^ ; split-GAL4-AD/+; split-GAL4-DBD/ UAS unc84 2xGFP in attP2 | 2A | anti-GFP, anti-Brp  Slowfade mounting  20x objective  Maximum Intensity projections through entire brain |
| pBPhsFlp2::PEST in attP3/w^1118^; split-GAL4-AD/+; split-GAL4-DBD/ pJFRC201-10XUAS-FRT>STOP>FRT-myr::smGFP-HA in VK0005, pJFRC240-10X-UAS-FRT>STOP>FRT-myr::smGFP-V5-THS-10XUAS-FRT>STOP>FRT-myr::smGFP-FLAG in su(Hw)attP1 | S2B (SS03656, SS00307, SS00308, SS02594) | anti-HA,anti-V5, anti-FLAG, anti-Brp  DPX mounting  63x objective  resampled views generated with Vaa3D |
| R57C10-Flp2::PEST in in attP18/ w^1118^; split-GAL4-AD/+; split-GAL4-DBD/ pJFRC210-10XUAS-FRT>STOP>FRT-myr::smGFP-OLLAS in attP2, pJFRC201-10XUAS-FRT>STOP>FRT-myr::smGFP-HA in VK0005, pJFRC240-10X-UAS-FRT>STOP>FRT-myr::smGFP-V5-THS-10XUAS-FRT>STOP>FRT-myr::smGFP-FLAG in su(Hw)attP1 | S2B (SS00321, SS00327) | anti-HA,anti-V5, anti-FLAG, anti-Brp  DPX mounting  63x objective  resampled views generated with Vaa3D |
| pJFRC19-13XLexAop2-IVS-myr::GFP in su(Hw)attP8, pJFRC21-10XUAS-IVS-mCD8::RFP in attP3/ w^1118^; split-GAL4-AD / PBac{fkh-GFP.FPTB}VK00037 ; split-GAL4-DBD /+ | 3K, S4H | anti-dsRed (recognizes RFP), anti-GFP  Slowfade mounting  40x or 63x objective  Single confocal sections |
| pJFRC19-13XLexAop2-IVS-myr::GFP in su(Hw)attP8, pJFRC21-10XUAS-IVS-mCD8::RFP in attP3/ w^1118^; split-GAL4-AD / PBac{Ets65A-GFP.FLAG}VK00037; split-GAL4-DBD /+ | S4I | anti-dsRed (recognizes RFP), anti-GFP  Slowfade mounting (DPX for C2 and C3 panels)  63x objective  Single confocal sections |
| pJFRC19-13XLexAop2-IVS-myr::GFP in su(Hw)attP8, pJFRC21-10XUAS-IVS-mCD8::RFP in attP3/ w^1118^; split-GAL4-AD / Mi{PT-GFSTF.0}TfAP-2^MI04611-GFSTF.0^; split-GAL4-DBD /+ | 4F,G | Native fluorescence (GFP, RFP)  Slowfade mounting  40x objective  Single confocal sections |
| y^1^ w^*^; Mi{PT-GFSTF.1}klg^MI02135-GFSTF.1^/TM6C, Sb[1] Tb[1] | 4I | anti-GFP, anti-CadN  Slowfade mounting  40x objective  Single confocal sections |
| pJFRC19-13XLexAop2-IVS-myr::GFP in su(Hw)attP8, pJFRC21-10XUAS-IVS-mCD8::RFP in attP3/ w^1118^; split-GAL4-AD / Mi{PT-GFSTF.0} Vmat^MI07680-GFSTF.0^; split-GAL4-DBD /+ | 5F | anti-dsRed (recognizes RFP), anti-GFP  Slowfade mounting  40x objective  Single confocal section |
| pJFRC19-13XLexAop2-IVS-myr::GFP in su(Hw)attP8, pJFRC21-10XUAS-IVS-mCD8::RFP in attP3/ w^1118^; split-GAL4-AD / Mi{PT-GFSTF.0}Nos^MI09718-GFSTF.0^; split-GAL4-DBD /+ | 5G | anti-dsRed (recognizes RFP), anti-GFP  Slowfade mounting  40x objective  Single confocal sections |
| w^1118^; PBac{y[+mDint2] w[+mC]=Ets65A-GFP.FLAG}VK00037 | 5H | anti-AstA (shown), anti-GFP (not shown)  Slowfade mounting  40x objective  Single confocal section |
| y^1^ w^*^ / w^1118^;; Mi{Trojan-GAL4.1}Oamb^MI12417-TG4.1^/ pJFRC12-10XUAS-IVS-myr::GFP in attP2 | 6B | anti-GFP, anti-repo  Slowfade mounting  40x objective  Single confocal section |
| y^1^ w^*^; Mi{PT-GFSTF.2}GluClalpha^MI02890-GFSTF.2^/TM6C, Sb[1] Tb[1] | 6C | anti-GFP, anti-repo  Slowfade mounting  40x objective  Single confocal section |
| pJFRC19-13XLexAop2-IVS-myr::GFP in su(Hw)attP8, pJFRC21-10XUAS-IVS-mCD8::RFP in attP3/ w^1118^; R47G08-LexAp65 in attP40 (Dm12)/ Lim3-GAL4 ^MI03817- Trojan-GAL4.1^ | S6C | Native fluorescence (GFP, RFP)  Slowfade mounting  40x objective  Single confocal section |
| pJFRC19-13XLexAop2-IVS-myr::GFP in su(Hw)attP8, pJFRC21-10XUAS-IVS-mCD8::RFP in attP3/ w^1118^; R65H03-LexAp65 in attP40 / Lim3-GAL4 ^MI03817- Trojan-GAL4.1^ | S6D | Native fluorescence (GFP, RFP)  Slowfade mounting  40x objective  Single confocal section |
| w^*^ P{y[+t7.7] w[+mC]=sens-FLPG5.C}attP18/ w^1118^; +/CyO; sens^Ly-1^/ TI{TI}VAChT^FRT-STOP-FRT.HA^ | 7A | anti-HA, native fluorescence (RFP in photoreceptors; present in sens-FLP clones due to perdurance)  Slowfade mounting  63x objective  Maximum Intensity projections through part of a confocal stack |
