## Supplemental Tables for "A genetic, genomic, and computational resource for exploring neural circuit function": TableS6.docx

**Homogenization buffer:**

Prepare a 500 – 1000ml stock of the following base buffer and store at 4°C.

mSodium acetate ph8.5

2.5mM MgCl_2_

250mM Sucrose

Add the following components to complete the homogenization buffer just before use (5mls/isolation).

| **Component** | **Stock**  **concentration** | **Amount to add to**  **5mls buffer** | **Final**  **concentration** |
| --- | --- | --- | --- |
| NP40 | 10% | 250ul | 0.5% |
| anti-GFP antibody | 0.2mg/ml | 10ul | 0.4ug/ml |
| spermidine | 1M | 3ul | 0.6mM |
| spermine | 1M | 1ul | 0.2mM |
| DTT | 1M | 5ul | 1mM |
| Protease inhibitor | Dissolve 1 tab in 1ml H2O | 100ul | 1X |
| Torula RNA | 10mg/ml | 250ul | 0.5mg/ml |
| Carboxyl beads | 30mg/ml | 100ul | 0.6mg/ml |

**Binding buffer:**

Prepare a 500 – 1000ml stock of the following base buffer and store at 4°C.

500mM Sodium acetate ph8.5

250mM Sucrose

6mM EGTA

6mM EDTA

Add the following components to complete the binding buffer just before use (5mls/isolation).

| **Component** | **Stock**  **concentration** | **Amount to add to**  **5mls buffer** | **Final**  **concentration** |
| --- | --- | --- | --- |
| spermidine | 1M | 3ul | 0.6mM |
| spermine | 1M | 1ul | 0.2mM |
| DTT | 1M | 5ul | 1mM |
| Protease inhibitor | Dissolve 1 tab in 1ml H2O | 100ul | 1X |
| Torula RNA | 10mg/ml | 125ul | 0.25mg/ml |
| Protein A beads | 30mg/ml | 30ul | 0.18mg/ml |

**Wash buffer:**

Prepare a 500 – 1000ml stock of the following base buffer and store at 4°C.

250mM Sodium acetate ph8.5

250mM Sucrose

Add the following components to complete the binding buffer just before use (10mls/isolation).

| **Component** | **Stock**  **concentration** | **Amount to add to**  **10mls buffer** | **Final**  **concentration** |
| --- | --- | --- | --- |
| NP40 | 10% | 500ul | 0.5% |

**Release buffer:**

Prepare 10mls of a 10X stock of the base buffer and store at 4 degrees C.

100mM Tris pH7.5

25mM MgCl_2_

5mM CaCl_2_

2.5mM Sucrose

Prepare 1ml of 1X Release buffer for both the trituration and post-release wash steps for each sample.

| **Component** | **Stock**  **concentration** | **Amount to add to**  **10mls buffer** | **Final**  **concentration** |
| --- | --- | --- | --- |
| Release buffer | 10X | 100ul | 1X |
| NP40 | 10% | 50ul | 0.5% |
| H2O |  | 850ul |  |

Just before use prepare the following 50ul reaction mix for a single bead release.

| **Component** | **Amount to add** |
| --- | --- |
| 10X Release buffer | 5ul |
| H2O | 31.5ul |
| RNAsin | 1ul |
| Torula RNA | 5ul |
| DNAseI | 1ul |
| IdeZ protease | 4ul |
| 10% NP40 | 2.5ul |
